# Carbohydrates complement high protein diets to maximize the growth of an actively hunting predator

**DOI:** 10.1101/2022.04.12.487891

**Authors:** Will D. Wiggins, Shawn M. Wilder

## Abstract

In nature, food is often variable in composition and availability. As a consequence, predators may need to seek non-prey food sources. Some predators are known to feed on nectar when food is limited. Nectar and other carbohydrate resources could also be beneficial when prey are more abundant if it helps predators balance protein-biased diets. We tested if an actively hunting predator, the jumping spider, *Phidippus audax*, benefited from liquid carbohydrates when prey were not limited. We also tested if the benefit of carbohydrates varied with the nutrient content of prey (i.e., from protein to lipid-biased). Spiders were reared on one of six live prey, *Drosophila melanogaster*, treatments that ranged from high protein to high lipid. Half of the spiders were given access to a 20% sucrose solution. After two months, we measured spider mass, cephalothorax width, instar duration, percent body fat, survival, and estimated number of prey eaten. Spiders reared on high protein diets with carbohydrates were larger and heavier than spiders on other treatments. Access to carbohydrates also increased percent body fat and survival across prey treatments. Our results suggest that carbohydrates may be a valuable component of spider diets, especially when prey have high protein and low lipid content as is commonly observed in prey in the field. Our results highlight the importance of diet balancing for predators, and that liquid carbohydrates can be an important nutrient to supplement a diet of prey rather than just being an energy supplement during periods of starvation.

**Significance statement:** Protein and lipid are thought to be the primary nutrients used by predators, including spiders. Yet, some spiders have been observed feeding on carbohydrate-rich nectar from flowers. We tested if the addition of carbohydrates to a high protein or high lipid diet affected the growth of the North American jumping spider, *Phidippus audax*. Spiders grew largest on high protein diets with carbohydrates, demonstrating that plant-based foods rich in carbohydrates can be important for some predators.

## Introduction

The quantity and quality of food consumed has large effects on the growth, reproduction, and survival of animals. Food is often limiting, especially for some predators, and individual food items can be nutritionally imbalanced or biased (Sterner & Elser 2002). Hence, animals can spend significant time foraging and often need to seek multiple food sources to balance their diet (Simpson & Raubenheimer 2012). When given a choice, many animals are capable of selecting food items that maximizes fitness by selecting foods that are nutritionally complementary (Lee et al. 2008, Maklakov et al. 2008). However, in nature free choice among nutritionally complementary foods may not always be possible and, hence, it is important to understand how limited or nutritionally-biased diets affect growth and survival.

Macronutrients are used for two broad functions: to fuel metabolism, and to provide building materials for body structures. Carbohydrates and lipid are often the primary source of energy while protein is often used to build new tissue. Protein can also be used as a source of energy (e.g., via gluconeogenesis) if other sources of energy are limited in the diet (Myers and Klasing 1999, Eisert 2011) or as a specialized source of energy (e.g., proline use by some flying insects; Klowden 2007). However, use of protein as an energy source is likely less efficient than other sources of energy (i.e., carbohydrates and lipid), and may produce harmful nitrogenous metabolic byproducts (Klowden 2007). While some animals can substitute macronutrients, like using protein as an energy source, not all animals are capable of macronutrient substitution. For example, cats have a limited ability to digest carbohydrates and cannot use this macronutrient as a substitute for low levels of lipid in their diet (MacDonald et al. 1984, Hewson-Hughes et al. 2011). In another example, carbohydrates are a non-substitutable resource for fire ants, with colonies able to increase both brood production and the number of worker when carbohydrates are available compared to colonies with only *ad libitum* lipid and protein from prey (Wilder et al. 2011). Understanding whether or how well macronutrients can be substituted, and/or complement one another, is critical for predicting the consequences of food limitation and nutrient imbalances for individuals and populations in nature (Simpson et al. 2006).

One factor that has a key influence on the primary nutrients used by animals is the trophic level at which they feed. Herbivores primarily ingest carbohydrate and protein, which are the bulk of the macronutrients in plants, and often consume limited amounts of lipid in their diet (e.g., some seeds). On the other hand, carnivores typically consume diets high in protein and lipid but low in carbohydrate (Russel et al. 2003). While the diets of herbivores and carnivores typically differ, both groups can engage in omnivory, with herbivores consuming animal tissue (White 2011) and carnivores consuming plant material (Wäckers et al. 2005). These deviations from what are considered typical diets for animals within a given trophic level are often thought to be an extreme response to starvation or as part of a coevolved food-for-protection mutualisms (Wäckers et al. 2005, White 2011, Ballova et al. 2015). Yet, recent research suggests that not all animals follow traditional views of being strictly carnivorous or herbivorous and that many, if not most animals, may be somewhat omnivorous (see the extensive review of Coll and Guershon 2002). There is a breadth of literature covering food-for-protection mutualisms in which plants or honeydew-producing hemipterans provide carbohydrates to carnivorous insects (e.g., ants and wasps) in exchange for protection (Wäckers et al. 2005), and even some literature on spiders feeding on plants (see Nyffeler et al. 2016). However, further work is needed to resolve the contribution of non-traditional food sources to the diets of animals, especially focusing on the role of plant-based foods in carnivore diets.

Spiders provide an interesting system for testing the role of plant-based foods for carnivores. Nearly all spiders are obligate carnivores (but see Meehan et al 2009). Yet, up to 30 species have been observed feeding on floral or extrafloral nectar in nature, including species in the wandering spider families Miturgidae, Thomisidae, Anyphaenidae, Corinnidae, and Salticidae (Jackson et al. 2001). In the miturgid spider, *Cheiracanthium inclusum*, supplementing the diet of food-limited individuals with nectar allows them to achieve growth and reproductive rates comparable to individuals fed higher quantities of prey (Taylor and Pfannenstiel 2009). Similar results have been found in the crab spider *Ebrechtella tricuspidata* (Wu et al. 2011), with honey (acting as simulated nectar) increasing survival and decreasing development time. Hence, some spiders appear to be able to use the carbohydrates in nectar to compensate for a lack of overall food availability. However, it remains unclear whether carbohydrates are only a source of nutrition during starvation or if carbohydrates contribute to growth of spiders when prey are more abundant. Furthermore, recent research suggests that the lipid and protein content of prey can have large effects on the growth (Wiggins and Wilder 2018), survival (Jensen et al. 2010) and reproduction (Lomborg and Toft 2009) of spiders and other predators. Macronutrient balance is integral to proper function, yet the availability of these key macronutrients can vary widely in prey (Wilder et al. 2013). As such, carbohydrates would be predicted to be more beneficial to spiders fed protein-biased prey as the carbohydrates could provide a source of energy to substitute for the low lipid content of prey (Noreika et al. 2016). Although, context-dependency in the benefit of carbohydrates for spiders remains to be tested.

The overall goal of this study was to test if a common plant-based food, liquid carbohydrates, benefitted the growth of an obligate predator when prey were not limited. Furthermore, we tested if the benefit of carbohydrates varied with the nutritional content of the prey, which can vary widely (Wilder et al. 2013, Wiggins and Wilder 2018). Specifically, we provided the jumping spider *Phidippus audax* with one of six diets of live prey (*Drosophila melanogaster*) that varied in their nutrient content (i.e., ranging from low to high lipid:protein) either with or without access to supplemental carbohydrates. We hypothesized that the addition of carbohydrates would have more of a benefit to the growth of spiders fed protein-biased prey than spiders fed lipid-biased prey.

## Methods

### Spider Maintenance

Spiders used in the laboratory experiments were first generation individuals whose parents were collected as penultimates during October-November 2015 from the old-field community surrounding Sooner Lake Dam, Pawnee Co., Oklahoma. The parent spiders were fed 1-2 appropriately sized crickets, *Acheta domesticus*, and watered twice a week. Parent spiders were paired for mating in mid-December. Spiderlings hatched in mid-January and were raised with their mother until they underwent their first molt. Twelve spiderlings from each females’ first clutch (n = 27) were separated into individual containers and given an alpha-numeric identification code (n = 324). Spiderlings were housed in Carolina Biological (Carolina Biological Supply Co., Burlington, NC) fly vials (3.3 cm diameter × 11 cm tall) with 2 cm of Plaster of Paris in the bottom to retain moisture and stoppered with sponge stoppers. The sponge stoppers had a small hole cut in the center. A translucent polypropylene drinking straw stuffed with cotton was inserted into the hole (diameter 5 mm x length 50 mm). The spiders were kept on a 14:10 hour light/dark cycle at a constant temperature of 26 °C.

### Prey Nutrient Treatments

We manipulated the macronutrient content of live prey items, wild type *Drosophila melanogaster*, by raising the flies on media with different nutrient content that allowed us to create six treatments of flies with particular ratios of lipid:protein (as in Jensen et al. 2011). As established in a previous study (Jensen et al. 2011) all the prey diets used Carolina Biological fly media (potato flake) as the base. Casein (milk powder) was added to the media to increase the protein content of the resulting flies or sucrose was added to increase the lipid content of the flies. Casein treatment ratios were 2:3, 1:4, or 1:9 casein to Carolina by mass. Sucrose treatments were either 1:2 or 1:4 sucrose to Carolina by mass, and one treatment was Carolina fly media with no supplemental nutrients. These treatments produce flies with a wide range of lipid:protein content (Table 1) from highly protein-biased (i.e. 2:3 casein:Carolina) to highly lipid-biased flies (i.e. 1:2 sucrose:Carolina) (see Jensen et al. 2011, Wiggins & Wilder 2018). Spiders were each fed four flies twice a week. Spiders were allowed twenty-four hours to consume the flies. After twenty-four hours, the flies were counted and released to give us an estimate of how many prey items were consumed. We are using the term estimate because we did not see each fly get consumed and cannot rule out natural deaths or kills without feeding.

**Table 1.** Fly diet effect on fly body composition and mass ± SD

| Fly diet | Fly body composition (lipid : protein) | Fly mass (mg) |
| --- | --- | --- |
| Casein 2:3 | 0.09 | $0.31 \pm 0.05$ |
| Casein 1:4 | 0.12 | $0.34 \pm 0.02$ |
| Casein 1:9 | 0.16 | $0.32 \pm 0.05$ |
| Carolina | 0.27 | $0.34 \pm 0.03$ |
| Sucrose 1:4 | 0.37 | $0.29 \pm 0.04$ |
| Sucrose 1:2 | 0.43 | $0.28 \pm 0.03$ |

To determine the role of carbohydrates in spider nutrition, we gave spiders in each diet treatment access to either a 20% sucrose solution (20 g of sucrose into 100ml of water, colored with 5 drops of red food coloring) or tap water (100 ml colored with 5 drops of red food coloring). The solutions were presented via the translucent drinking straws stuffed with cotton. We used red food coloring in both solutions to verify that spiders were drinking the solutions. When spiders drank the food coloring it turned their excreta a red-reddish brown color. Also, the abdomens of the spiders that drank the red solution appeared pinkish. All spiders were given access to water without food coloring twice a week via a light misting when flies were removed.

We alternated between providing spiders with liquid solutions and flies. The two were never provided at the same time, as not to allow the flies to drink the sucrose solutions and change their macronutrient contents. The straws with water or sugar water were inserted for seventy-two hours, until the next feeding of flies.

### Measurement of Growth

Spider growth was calculated using multiple measures. First, spiders were weighed on a scale to the nearest 0.01 mg at two months. The first molt following this mass measurement was recorded for date. Molts were collected and measured for a fixed body size, carapace width at the posterior lateral eyes (PLE). We used carapace width alone because the other common size measurement, patella/tibia length, is highly correlated with carapace width (Wiggins and Wilder 2018). Photos were taken of the molts and a micrometer slide using a camera attached to a dissecting microscope and measured with ImageJ software (Rasband 2016) to 0.001 mm. These measurements provide an accurate measure of the size of the spider’s body at weighing without undue stress or sacrificing the animal. Some molts were damaged before or during the weighing, slightly decreasing total sample size.

### Flies Consumed

Twenty-four hours after feeding, surviving flies were counted and released. We are able to estimate the number of flies eaten by taking the total flies fed (4 flies twice a week for two months) and subtracting the number of flies released.

### Spider Lipid Content

We analyzed a subset spiders from each treatment (n = 76) for lipid content using a gravimetric protocol. Spiders were dried in an oven for 24 hours at 60 C° and weighed to the nearest 0.01 mg. Following the initial dry mass measurement, spiders were washed in chloroform for two consecutive twenty-four hour baths, with the chloroform being changed between baths. The lipids from the spiders were solubilized within the chloroform. After the second bath, spiders were once again dried in an oven for 24 hours at 60 C° and reweighed to the nearest 0.01 mg to obtain the final lean (i.e., lipid-free) mass.

### Statistical Analysis

We used Generalized Additive Models (Woods 2006) to test for effects of prey nutrient content, sugar access, and their interaction on spider growth. Generalized Additive Models are similar to Generalized Linear Models without the assumption of linearity. This allows the response to take a nonlinear form. The GAMs were run in R (R Core team 2014). JMP (SAS Inc. 2016) was also used to analyze survival of the spiders via a parametric survival model with a Weibull distribution analyzing the effects of prey nutrients, supplemental carbohydrates, and the interaction between prey nutrients and carbohydrates on spider survival. JMP was also used for all post-hoc analyses.

## Results

### Mass

There were significant main effects of both carbohydrates (*f* _1,302_ = 65.96, *p* < 0.01), and prey nutrients (*f* _5,298_ = 76.86, *p* < 0.001), as well as an interaction between prey nutrient content and carbohydrates on spider mass (*f* _6,297_ = 24.11, *p* < 0.001) (Figure 1a). We further explored this interaction with post hoc tests. For spiders fed the three highest lipid prey (i.e., highest lipid:protein), there was no significant difference between mass of individuals with or without carbohydrates (Figure 1a). However, the spiders fed the three highest protein prey treatments were significantly heavier when supplemented with carbohydrates (Figure 1a). The difference between spider mass with and without carbohydrates increased as the prey treatments became more protein biased (*f* _1,6_ = 10.86, *p* = 0.03).

**Figure 1.**
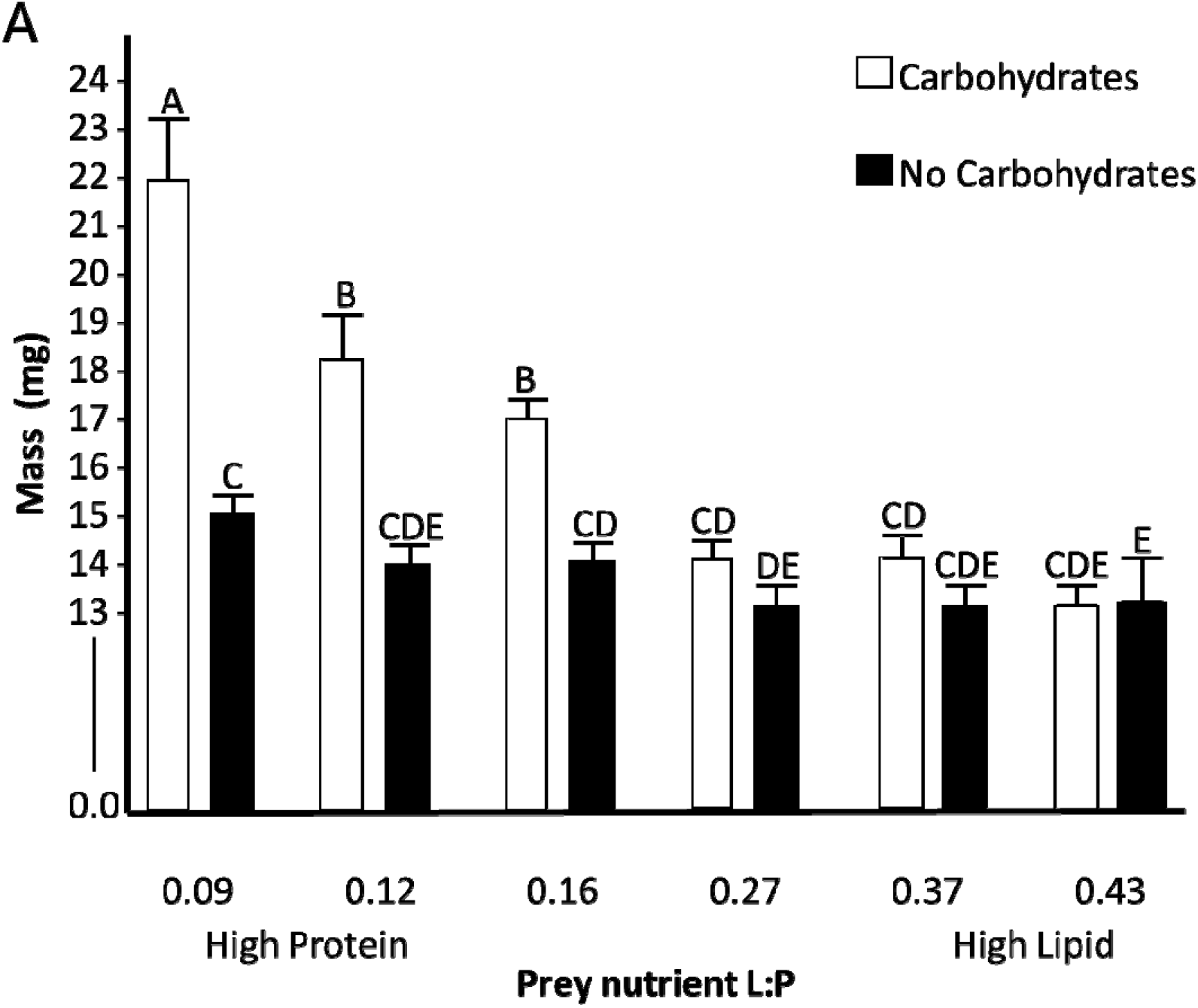
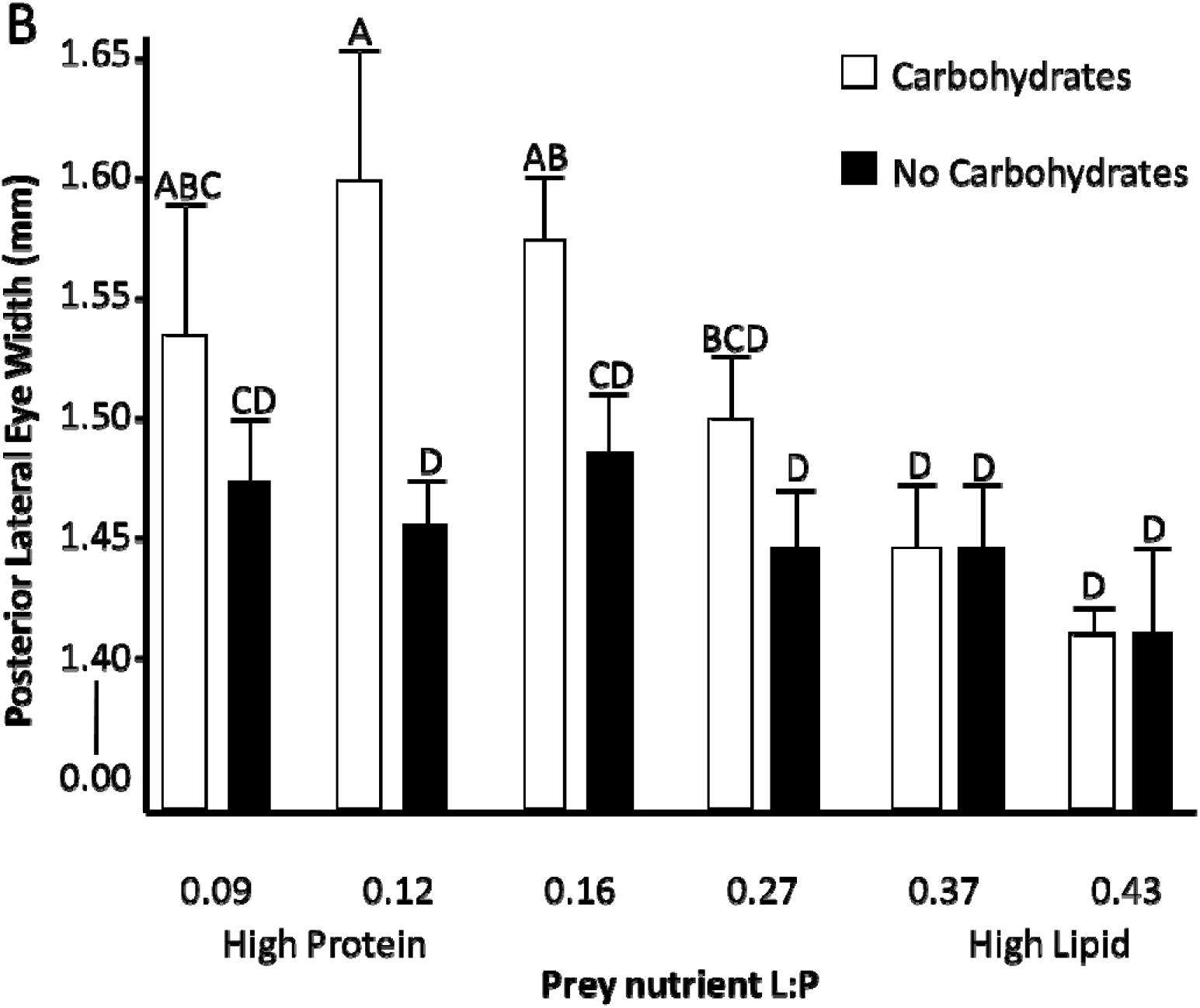
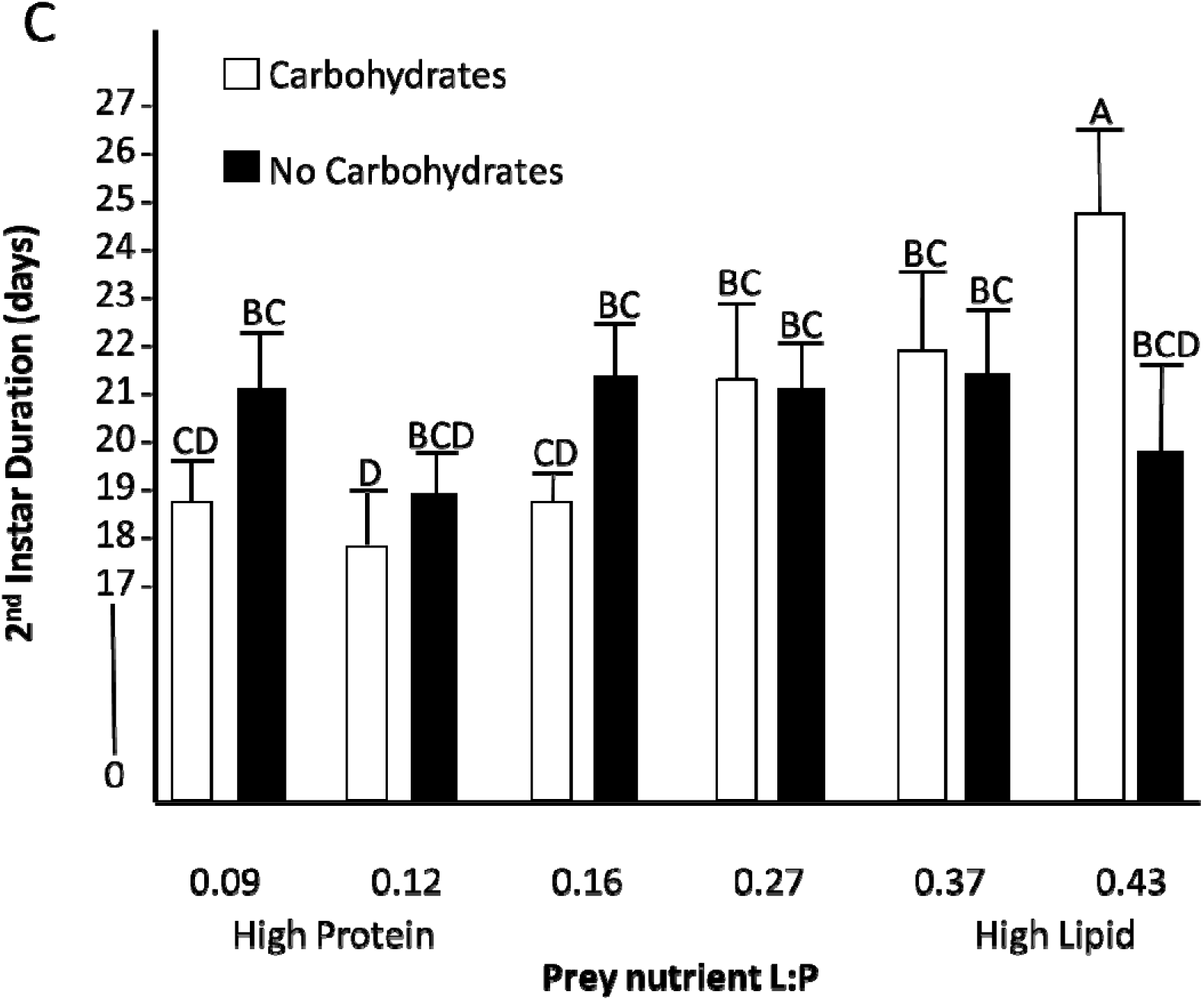
Growth metrics of spiders after being fed flies, one of six different prey nutrient ratios, ranging from high protein to high lipid with and without an available carbohydrate source at two months, with post hoc assignments and standard error. a) spider live mass b) posterior lateral eye width c) second instar duration

### Body Size

There were significant effects of carbohydrates (*f*_1,278_ =15.50, *p* < 0.001), prey nutrients *f*_6,273_ = 21.84, *p* < 0.001) and their interaction *f*_7,272_ = 6.61, *p* = 0.01) on spider body size, measured as carapace width at PLE (Figure 1b). Post hoc tests showed that on the high protein prey treatments, spiders provided carbohydrates had significantly wider carapaces than those without carbohydrates. There were no differences in carapace width of spiders fed high lipid prey regardless of carbohydrates presence.

### Instar Duration

We measured instar duration for both the 2^nd^ and 3^rd^ instars. We found a significant effect of prey nutrient content for the second instar duration (n = 279, *f* _5,273_ = 17.03, *p* < 0.001) and a significant interaction between prey nutrients and supplemental carbohydrates *f* _7,271_ = 13.68, *p* < 0.001). The main effect of carbohydrates was not significant (*f* _1,277_ = 0.12, *p* = 0.73). Post hoc tests indicated that spiders fed high lipid prey items with available carbohydrates took significantly longer to molt than all other treatment groups. To further explore the interaction effect, we conducted linear regressions of instar duration and prey lipid:protein separately for carbohydrate and no carbohydrate treatments. Linear regression of data from only carbohydrate supplemented spiders showed that individuals molted sooner when fed prey with higher protein content (n = 138, *f* _5,132_ = 33.05, *p* < 0.001). However, for spiders not provided carbohydrates, there was no effect of prey nutrient content on second instar duration (n = 141, *f* _5,135_ = 0.09, *p* = 0.77) (Figure 1c). We took the same measurements for the 3^rd^ instar duration and found that there were no longer any significant effects of prey nutrients (n = 109, *f* _5,103_ = 1.86, *p* = 0.18), supplemental carbohydrates (*f* _1,107_ = 2.20, p = 0.14), nor the interaction (*f* _7,101_ = 0.008, *p* = 0.78) on instar duration.

### Survival

Of the starting spiders, 199 individuals survived and 108 died (sample size is less than total spiders in the experiment due to some escapes n = 17). An effects likelihood ratio test revealed that prey nutrient content did not significantly affect survival (χ^2^ = 0.08, df = 1, *p* = 0.78). However, the presence or absence of supplemental carbohydrates did significantly affect survival (χ^2^= 25.97, df = 1, *p* < 0.001), with individuals fed carbohydrates having higher survival. The interaction between prey nutrients and carbohydrates was near significant (χ^2^ = 3.60, df = 1, *p* = 0.06), but no clear conclusions could be drawn (Figure S1).

#### Spider Lipid Content

We measured the body fat content of a subset of spiders that survived across all nutrient treatments. We found that the presence or absence of supplemental carbohydrates was the only factor that affected body fat content (n =76, *f* _1,74_ = 21.78, *p* < 0.001), with spiders fed carbohydrates having higher body fat content than spiders not provided carbohydrates (Figure 2). Prey nutrients and the interaction were non-significant (*f* _5,70_ = 0.007, *p* = 0.93; *f* _7,68_ = 0.75, *p* = 0.39, respectively).

**Figure 2.**
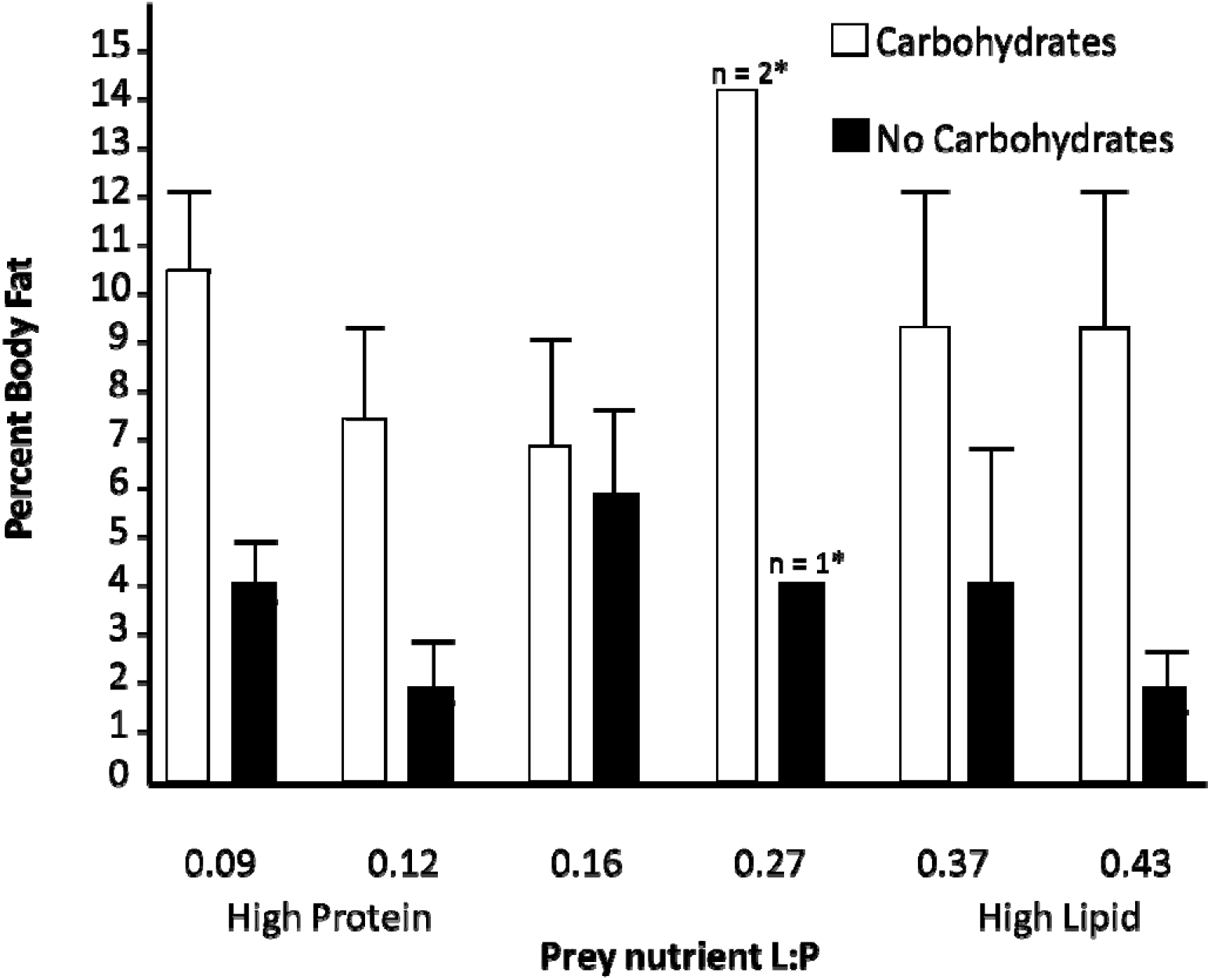
Percent body fat from surviving spiders fed one of six different prey nutrient ratios with and without an available carbohydrate source. * Denotes groups with too few samples for standard deviation.

#### Total Prey Consumption

There was a significant effect of prey nutrients (n = 301, *f* _5,295_ = 6.95, *p* = 0.009), carbohydrates *f* _5,299_ = 15.71, *p* < 0.001), and the interaction (*f* _5,293_ = 26.13, *p* < 0.001) on the number of flies consumed by spiders (Figure 3). Post hoc tests revealed that spiders fed high protein diets with carbohydrates ate fewer flies than spiders fed high protein diets with water (Figure 3).

**Figure 3.**
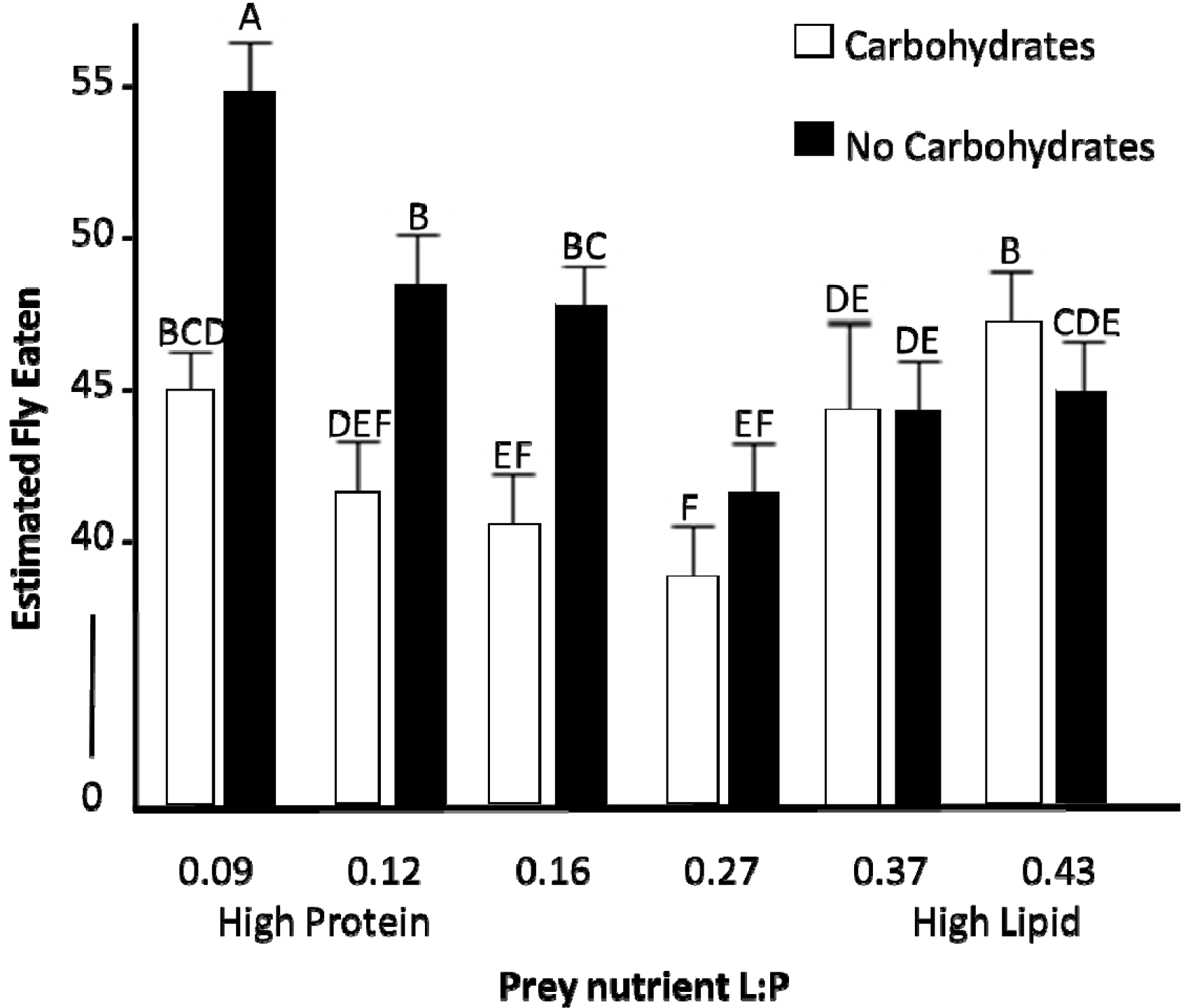
Estimated total flies consumed after 2 months of sustained feeding of eight flies a week across six different prey nutrient ratios with and without an available carbohydrate source. Post hoc assignments and standard error are shown above the mean, indicated by the bar.

## Discussion

Our results demonstrate that carbohydrates can be a valuable component of carnivore diets, especially when carnivores are feeding on high protein prey. Spiders fed the high protein prey and also provided carbohydrates were almost twice as heavy and had wider cephalothorax than spiders in all other treatments. Instar duration was longest for spiders fed carbohydrates and high lipid prey, putting them at a disadvantage but only during the second instar. The effect was no longer present in the third instar. Supplementation with carbohydrates also increased survival and percent body fat across all prey nutrient treatments. We had expected carbohydrates to increase growth, but the large benefit of carbohydrates for spider growth (i.e., carbohydrate fed spiders were > 40% heavier than non-carbohydrate fed spiders on the highest protein treatment) was not expected for such an obligate carnivore. Previous studies have found that spiders, especially actively hunting species, will consume nectar in nature and that nectar can increase survival and growth during periods of limited food availability or starvation (Taylor and Pfannenstiel 2009, Wu et al. 2011). Our results suggest that carbohydrates may be more than just an energy supplement during periods of starvation and that carbohydrates could be an important component of this actively hunting spider’s diet during development.

The high protein diet with carbohydrates may have provided the ideal situation for spiders: a large amount of protein to build new tissue and carbohydrates to provide a source of non-protein energy for growth. The high protein diet offered the highest amount of protein to invest in new tissues. However, the high protein prey lacked a readily metabolizable energy source. Protein can be catabolized for energy, but doing so is less efficient than extracting energy from carbohydrates or fat. Also, this metabolizes the protein so that it cannot be used to build tissue. The spiders on higher lipid diets also had a balance of protein and non-protein energy (lipid) but they had less overall protein and the addition of carbohydrates would only provide additional energy, which may not have been limiting on this diet. This is similar to what is observed in carabid beetles, where lipids and carbohydrates can be used interchangeably (Noreika et al. 2016). The present results suggest that the observations of spiders feeding on nectar in nature may be due to diet choice by the spiders, especially since prey are often protein-biased in nature (Wilder et al. 2013, Wiggins and Wilder 2018).

Spiders fed the high protein prey with no carbohydrates ate significantly more flies than spiders fed the high protein prey with carbohydrates. By feeding on more prey, spiders on the high protein treatment could have been either selectively extracting the limiting lipid from many prey (e.g., Mayntz et al. 2005) or consuming large amounts of protein to catabolize some of this protein for energy. Measurement of nutrients in the prey carcasses would have been needed to differentiate between these mechanisms. Regardless, the differences among treatments in fly consumption demonstrate that spiders are able to adjust their foraging behavior to compensate for variation in the nutritional composition of prey or available resources and its potential consequences for growth. Jensen et al. (2011) demonstrated similar compensatory feeding in *Pardosa prativaga* fed prey varying in lipid and protein content and found relatively few effects of diet on spider growth. Compensatory feeding in the absence of nectar could have important implications for understanding spatial and temporal variation in food web dynamics and how it may relate to the availability of floral resources.

In addition to providing more energy, there could be a difference in the digestibility of carbohydrates relative to the other major energy source, lipid. For example, studies of fire ants have shown that the addition of liquid carbohydrates (an artificial nectar substitute) increased colony growth even when insect prey, which contained significant amounts of lipid and protein, were available *ad libitum* (Wilder et al. 2011). Also, Toft and Nielsen (2017) have observed differences in carabid beetle metabolism of carbohydrates versus lipids and protein, and found carbohydrates best for replenishing fat reserves post-hibernation. It is possible that carbohydrates may, similarly, be more readily metabolized than lipids by spiders as well. Further work is needed on the metabolic costs of digesting (i.e., specific dynamic action, SDA) different foods and nutrients in spiders and other predators.

A previous study using similar fly diets with no carbohydrates found that juvenile *P. audax* grew largest on flies with the highest lipid content (Wiggins and Wilder 2018). The present study did not find similar results for the no carbohydrate treatments. Another interesting difference between the studies is that the spiders in the present study at two months of age were larger than spiders in the past study at four months of age (Wiggins and Wilder 2018). There are at least two potential explanations for the differences between the studies. First, the present study provided spiders with more readily available water in wet cotton, whereas the past study only periodically provided a spray of water droplets in containers. Water availability could interact with nutrient content of prey to affect spider growth (McCluney 2017). Second, the maintenance fly cultures used to inoculate our fly treatment cultures were raised in two different ways. In the past experiment, maintenance flies were cultured on only potato flake medium versus the present study where they were maintained on potato flake supplemented with ground dog food. While the flies fed to spiders had similar macronutrient content in both studies, there could have been transgenerational effects of past culture on different media that affected some unmeasured aspect of fly quality. The differences between the past and present studies suggest that while macronutrients can be important factors affecting prey quality, there may be other aspects of prey that can affect predator growth.

Across all diets, carbohydrates increased spider survival, with between 20% to 60% more spiders surviving when carbohydrates were present. This survival benefit is likely to be even higher in nature due to the benefit of carbohydrates for increasing lipid reserves. Increased lipid reserves would help spiders during food limitation, which can be often for some species (Wise 2006). Studies of a linyphiid spider suggest that they regularly experience periods of starvation of one week or more in nature (Bilde and Toft 1998). Studies of a wolf spider showed that wild-caught spiders had body condition not significantly different from lab-maintained spiders that were fed *ad libitum* and then completely deprived of food for three months (Wilder and Rypstra 2008). Lipid reserves are critical for surviving periods of starvation and, regardless of the prey on which they fed, spiders that consumed carbohydrates had higher lipid reserves than spiders that did not have access to carbohydrates.

These results demonstrate that carbohydrates can be an important component of spider diets. There are at least two potential ways that spiders may consume carbohydrates in nature: 1) by feeding on pollinators that have recently fed on nectar, and 2) by feeding directly from the plant, either through floral or extrafloral nectaries. It is likely that jumping spiders consume carbohydrates from both of these mechanisms. Jumping spiders and some other wandering spiders hunt prey on flowers, as flowers provide a hotspot of insect activity. Many of the insects captured on flowers will likely have nectar in their guts from recently feeding on other flowers. Some jumping spiders have also been observed feeding directly from nectaries (see Nyffeler et al. 2016). Given the potential benefits of hunting from flowers, it is surprising that jumping spiders are not more specialized for this behavior (e.g., coloration to blend into flowers). Although, this could be due to competition for these hunting locations with other predators that frequently hunt on flowers (e.g., praying mantids, crab spiders). Rather, for jumping spiders, flowers may serve as one of multiple feeding sites used as the spiders actively move through their habitat.

## Acknowledgements

We would like to thank OG&E for allowing us access to the Sooner Lake Dam for the collection of paternal spiders. We would also like to thank Jodie Wiggins, Sarah Durant, Matt Lovern, and Jennifer Byrd-Craven for helpful comments on earlier versions, as well as two anonymous referees. The authors would also like to thank Oklahoma State University for the startup funds that funded this research. As well as the numerous undergraduates that helped maintain fly cultures and the lab, with special thanks to Sarah Bounds who assisted in weighing and measuring spiders.

## Data Availability

The datasets generated during and/or analyzed during the current study are available from the corresponding author on reasonable request.

